# Comment on ‘Response to comment on ‘Valid molecular dynamics simulations of human hemoglobin require a surprisingly large box size’’

**DOI:** 10.1101/812966

**Authors:** Vytautas Gapsys, Bert L. de Groot

## Abstract

We recently expressed three major concerns about a 2018 article of El Hage et al. about a claimed effect of the box size in molecular dynamics simulations of hemoglobin. In the response of the authors to our comment, none of these concerns have been addressed, yet the authors maintain their original conclusions. Here, we challenge those conclusions and provide additional data that reestablish our original concerns. In addition, we identified six additional flaws in the response from El Hage et al. as well as a number of technical concerns about the presented simulations and analyses. Taken together, we conclude that there is no basis to support the hypothesis of significant box size effects in MD simulations for the studied systems in the examined range.

## Introduction

A 2018 study reported on a claimed box size effect for hemoglobin T to R transitions in molecular dynamics simulations (1). In a recent comment, we challenged the conclusions of that work, in which we expressed three major concerns about that study (2). Briefly, we challenged the statistical basis of that study, identified normalization issues in the analysis, and argued that the reported transition times inform on kinetics of the transition rather than the thermodynamic difference between the endstates. In the response to our comment, our concerns were challenged, and the authors maintained their original conclusions on the simulation box size effect (3). Here, we respond to these challenges, argue why they do not address our original concerns, identify six additional flaws in the response, and provide additional data that challenge the claims of El Hage et al. We will start by discussing our original concerns and then follow with the additional issues introduced in the response to our comment.

## Results

### Statistics

First, we expressed our concern about the conclusions based on single replica (N=1) statistics presented by El Hage et al. (1), that remain unchanged in their response, as no additional data were added (3). We argue that, as molecular dynamics simulations behave as chaotic systems and therefore sample stochastically, a single simulation replica per simulation box size in which a transition is observed (or not) is not sufficient to support conclusions on box size effects on that transition. Rather, N=1 evidence is anecdotal in nature, as had been pointed out by one of the reviewers of the original paper (1) and in line with a recent investigation into the importance of mutiple replicas in MD simulations (4). Indeed, in our investigation, in which we applied an order of magnitude more statistics (N=10-20) (2), we found a large scatter of transition times for each studied box size. These were found to cover the full range of times in the El Hage et al. study, the differences between which had been attributed to an effect of the simulation box size. Rather than the claimed systematic trend of slower transitions for larger boxes as had been deduced by El Hage et al. from their N=1 statistics (1), the scatter observed from more repeats therefore indicates that the observed differences in transition times from single replicas are stochastic in nature.

Against this background, it is thus surprising that in their response to our comment, El Hage et al. now challenge our N=10-20 statistics, while maintaining their original N=1 based conclusions (3). How absurd this is can be appreciated from the fact that N=1 subsets can be readily extracted from our N=10-20 sets that would give rise to the exact opposite conclusion as that of El Hage et al. (i.e. a *faster* transition for larger box sizes rather than *slower*, see Fig. 1). This example shows that the presented N=1 basis of El Hage et al. does not suffice to support the conclusion of a box size effect on conformational transitions in hemoglobin.

**Figure 1:**
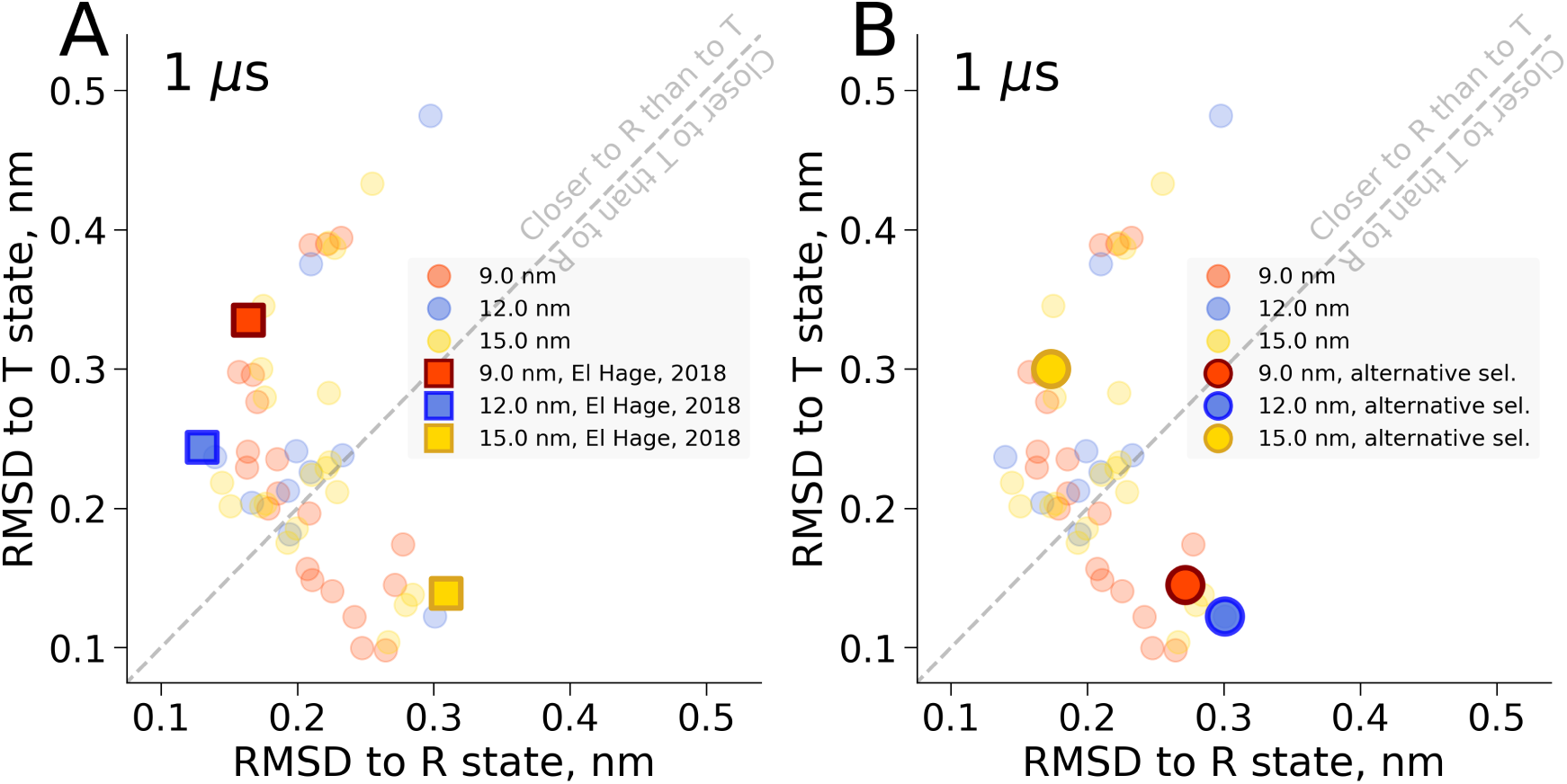
RMSD from crystallographic T and R state for simulation snapshots after 1 microsecond. Panel A highlights single simulations in which the 9 nm and 12 nm boxes make the T to R transition and the 15 nm box does not, as in El Hage et al. (1). Probability of observing this effect given the current sampling is 0.13 ± 0.06. Panel B shows an alternative selection of single simulations showing the opposite effect: the 15 nm box makes the transition, and the 9 nm and 12 nm boxes do not. Probability of this effect is 0.06 ± 0.05.

### Hydrophobic effect

In our second concern, we had identified that suggested differences in protein solvation for different box sizes, linked by the authors to the hydrophobic effect (1), are rather explained by normalization issues. Upon correction of these issues, rather than finding box size dependent differences in solvation for different simulation box sizes (1), we found highly similar (identical within error) solvation for all studied box sizes. Indeed, all previously reported differences by El Hage et al could instead be readily explained by an artificial, box size dependent normalization. We found this to hold true for solvent hydrogen bonds, diffusion constants as well as radial distribution function (RDF) (2).

Our initial RDF analysis rested on the extraction of a constantly sized analysis box from differently sized simulation boxes, to enforce consistent normalization. Potential issues with such a procedure were pointed out (3). We now implemented a spherical RDF normalization that does not rely on subsystem extraction from larger boxes. As can be seen in Fig. 2, the result of this radial normalization is consistent with our previous observation (2) and it confirms that no significant box size effects are observed in terms of protein solvation and therefore the hydrophobic effect.

**Figure 2:**
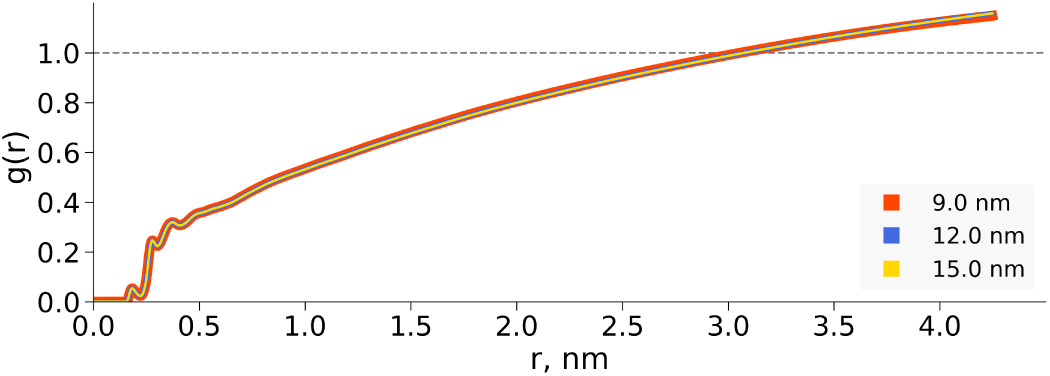
Water RDF for differently sized hemoglobin boxes, normalized to a sphere of 8.5 nm diameter.

### Kinetics rather than thermodynamics

Our third concern was regarding the conclusion of El Hage et al. about the thermodynamics of the T state relative to the R state as derived from a set of transition times for unidirectional T to R transition trajectories. A claim by El Hage et al. was made about the thermodynamic stabilization of hemoglobin’s T state in 15 nm box, based on the absence of a transition. We argued that for these stochastic barrier crossings, the transition rates primarily depend on the transition barrier, therefore not allowing any conclusions on the thermodynamic difference in stability between T and R. It is therefore surprising that in their response, El Hage et al. maintain their claim on thermodynamics based on a readout of unidirectional transition times, which are per definition kinetic.

In addition to these three original concerns, that remain unaddressed, in the following, we discuss several novel concerns that were introduced in the response of El Hage et al. (3).

### Number of transitions required for rate estimate

In their response (3), rather than addressing the insufficient N=1 basis in their original work, El Hage et al. instead question the N=10-20 statistics in our comment (2). They argue that converged reaction time distributions require hundreds to thousands of transitions. We would like to point out two flaws with this argumentation. First, if it were true that N=100-1000 were required for a converged rate estimate, then the N=1 transition times from their own data would clearly not be sufficient for a reliable rate estimate or indeed a conclusion about differences between one N=1 case and another. Hence, if anything, this would only underscore our original first concern that challenges the statistical basis of the data that underlies the box size hypothesis.

Second, we would like to point out that the N=100-1000 as claimed by El Hage et al. is a rather arbitrary threshold for the number of transitions required for a converged rate estimate. As would be expected (also see Fig. 3), the uncertainty in the rate estimate scales with 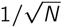, with N the number of times the transitions have been recorded. Therefore, the required size of N depends on the required level of uncertainty or confidence for the question at hand. A relatively small N, e.g. N=10, already suffices for a standard error of less than 50%, i.e. sufficient to detect rate *differences* of a factor 2 or more. Conversely, similar or identical rates will give rise to overlapping distributions of transition times, where the uncertainty in the rate estimate provides an upper bound for the difference in the underlying rates. The latter is exactly what we found for the hemoglobin data with N=10-20 (2): overlapping transition times for different box sizes indicating identical or highly similar transition rates for hemoglobin in differently sized simulation boxes. Bootstrap analysis allows testing the hypothesis put forward by El Hage et al. that the largest employed (15 nm) simulation box induces a significantly slowed down transition rate (3). As pointed out previously (2), based on the presented N=10-20 statistics, only a fraction of 0.0026 of the scenarios covered by bootstrap would be consistent with a moderate slowdown of a factor of 2 for the 15 nm box (according to the available statistics, a more substantial 10-fold slowdown is far less likely). In fact, we can use the same statistics to test the opposite hypothesis of faster transitions with larger simulation boxes. For that scenario, the computed probability of a 2-fold *faster* transition in the 15 nm box is 0.18. Thus, the employed N=10-20 in this case are sufficient to reject with a large confidence margin the hypothesis put forward by El Hage et al. that the largest employed (15 nm) simulation box induces a significantly slowed down transition rate. Increasing the number of trajectories to 100, as suggested by El Hage et al. as a minimal requirement (3), does not change the estimated rates significantly, see also Fig. 4.

**Figure 3:**
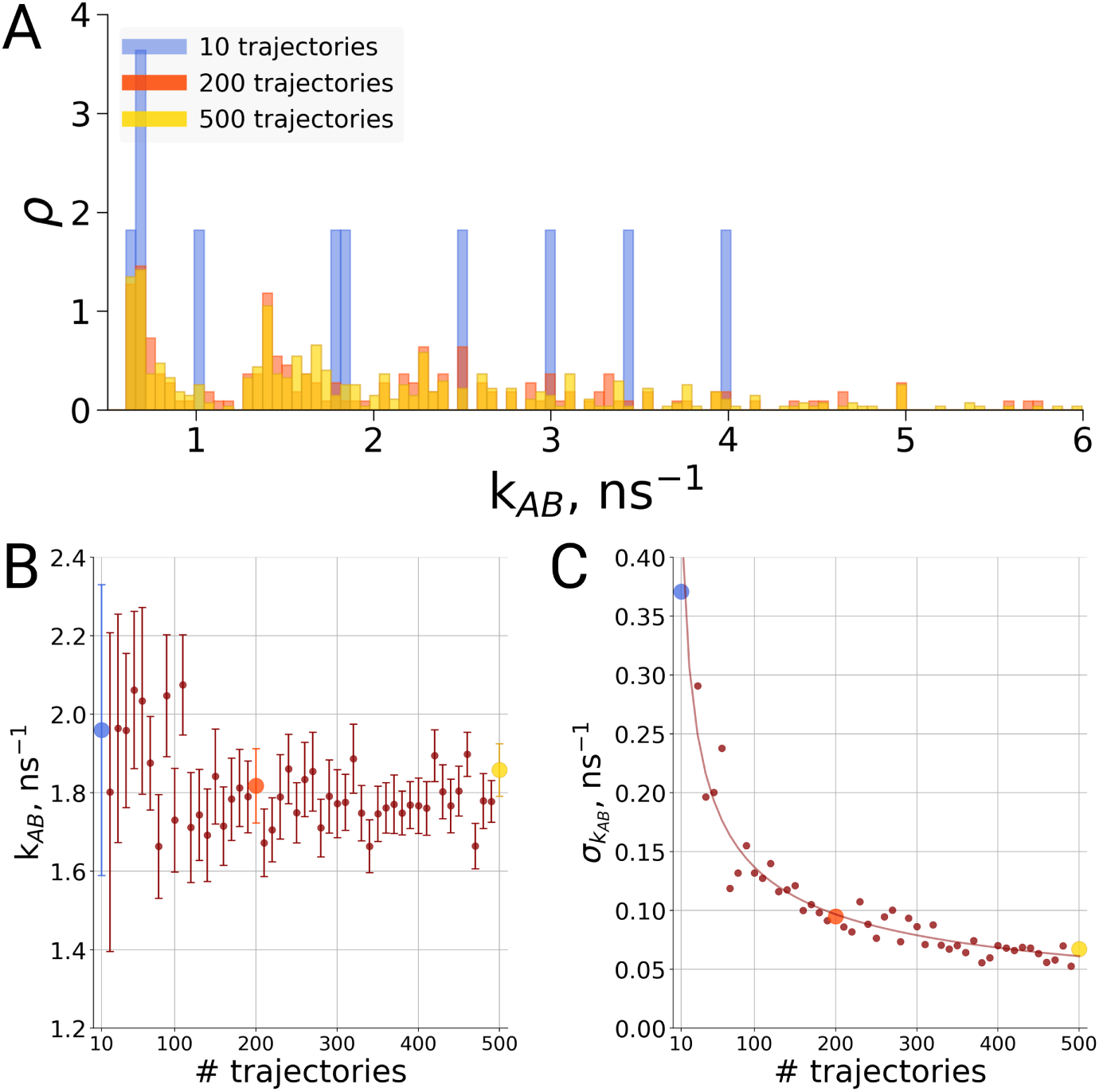
Rate estimate dependence on the number of transitions between the two main conformations of alanine dipeptide (backbone dihedral angle ψ is used as a reaction coordinate, the examined conformations are described in detail in (2)). Panel A shows a distribution of transition frequencies for sample sizes of 10, 200 and 500 trajectories. In B, the rate estimate is shown as a function of the number of trajectories and in C the associated uncertainty is plotted. In B and C the sample sizes of 10, 200 and 500 are highlighted with blue, orange and yellow spheres, respectively. The line in panel C depicts 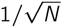, scaled to the least squares fit uncertainty.

**Figure 4:**
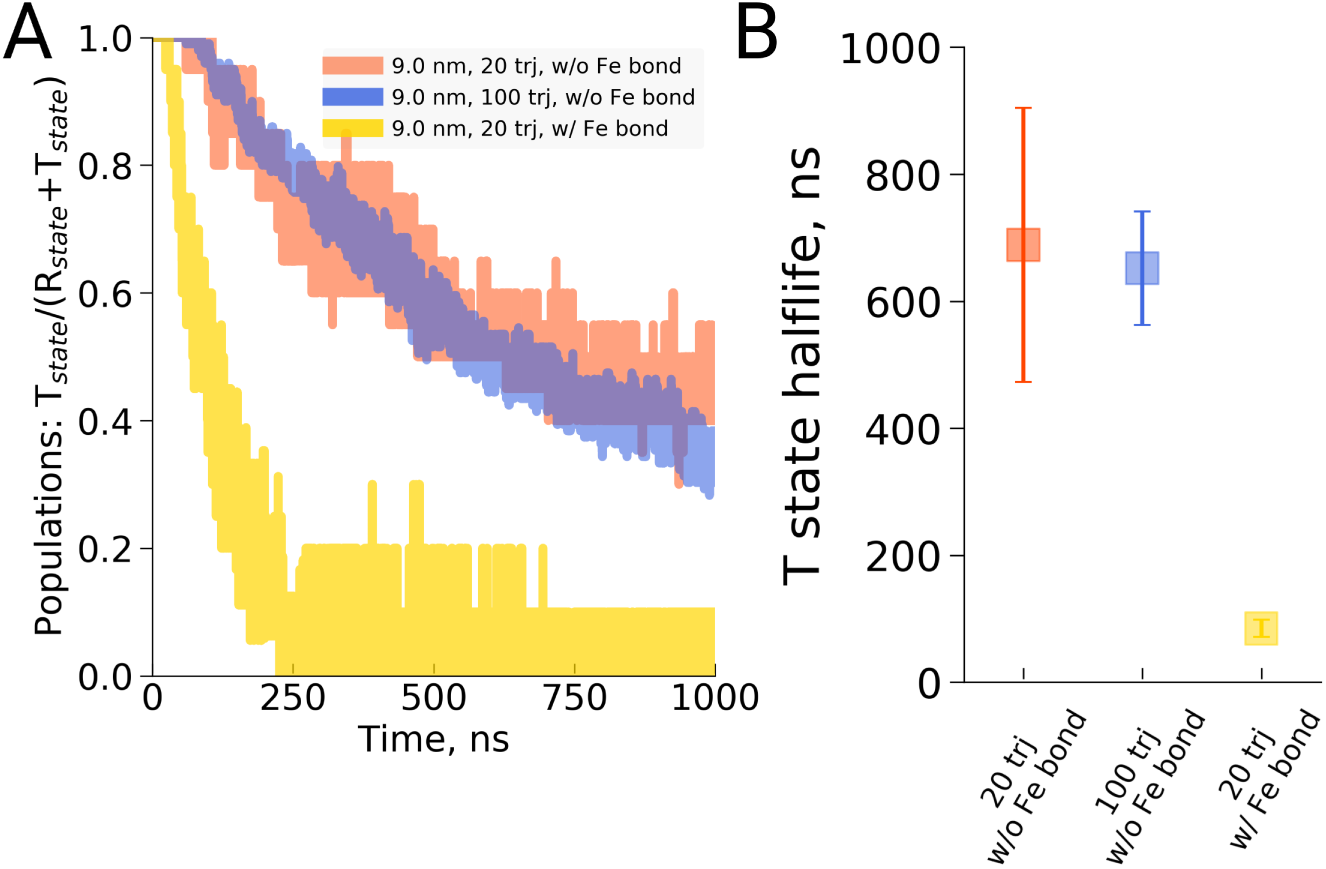
Hemoglobin T to R transition time statistics for different numbers of transition trajectories as well as for different bonding schemes between the iron and proximal histidine. Panel A shows the relative population of trajectories that remain in the T state as a function of time, whereas panel B shows the estimated mean transition times, together with their estimated uncertainty. 20 and 100 simulations with the iron-proximal histidine interaction modeled as a non-bonded interaction show consistent transition times, whereas the case in which the iron-proximal histidine interaction is modeled as a covalent bond shows a drastically reduced mean transition time.

### Logical flaw of misrepresenting claims

In a number of places in the response by El Hage et al. (3), assertions from our comment (2) are misrepresented leading to a logical flaw. As an example, the response contains the phrase: “Gapsys and de Groot have commented on this work but do not provide convincing evidence that the conclusions of El Hage et al., 2018 are incorrect.” What we had argued in our comment is that the conclusions by El Hage et al. (1) are not supported by the data. There is an important difference between “not supported” and “incorrect”, in particular concerning the “convincing evidence” that would be required to support the respective claim as well as on which side the burden of proof is (5). In our comment, we argue that based on the three expressed concerns, the data of El Hage et al. do not support the conclusion of a box size dependence. This implies that it is the responsibility of El Hage et al. to provide the evidence (e.g. in the form of additional data) to reach the point where the data is conclusive, or, alternatively, modify the conclusion. Neither was provided by El Hage et al.

Instead, the formulation of El Hage et al. attempts to shift the burden of proof to us, by misciting us as having claimed “that the conclusions of El Hage et al., 2018 are incorrect,” in our comment. Indeed, had we claimed that, it would have been our responsibility to provide data to conclusively show the absence of a box size effect. Rather, however, it is the responsibility of El Hage et al. to provide the data to support their conclusion of a box size effect, and not ours to provide evidence for the absence thereof.

Note that, in fact, we did provide a large amount of additional data that indidated that, if any, box size effects in MD simulations were rather small for the studied systems (2), but these findings are independent of the assertion that the conclusions by El Hage et al. (1) are not supported by their data, and thus do not shift the burden of proof to our side.

### Flawed assertion about thermodynamic stability

As a response to our concern that the reported transition times are about kinetics rather than thermodynamics, El Hage et al. in their response included the statement “A necessary but not sufficient criterion for thermodynamic stability is that the structure considered (here T(0)) does not decay on the time scale of the simulations (e.g., to R(0)).” This statement is incorrect. A state can be thermodynamically stable and yet show transitions to another state on short (simulation) time scales. An example of that is shown in the work of Daura et al. (6) that demonstrates the reversible folding of a β peptide. In simulations below the unfolding temperature, the folded helical state is the most stable conformation. Nevertheless, it shows frequent excursions to an unfolded conformation. Thermodynamic stability can thus not be read from only the decay rate to another state but must also consider back transitions. As we have shown in our comment, neither the forward nor the backward transitions show a significant trend as a function of box size, therefore not supporting the claim of El Hage et al. that the 15 nm box has a stabilizing effect on the T state of hemoglobin.

### Partial transition back to T state

In their response, El Hage et al. in Fig 3. added a simulation of a hemoglobin starting from the R state that showed a partial back transition to T, and write: “This is an additional indication that, thermodynamically, T(0) is favored over R(0) in the largest water box” (3). We argue that this assessment is flawed. First, it is once again again anecdotal evidence (N=1) that lacks the statistical significance to draw conclusions on thermodynamics (see first concern). Second, this test for possible partial back transitions was only carried out for the largest, 15 nm box. Lacking information from the other box sizes, it cannot be concluded that the effect is a feature of the largest box.

Our analysis over multiple replicas and different box sizes showed partial back transitions independently of the box size (Figure 1-figure supplement 1) (2). This further questions the assertion by El Hage et al. that one partial back-transition to T observed in the largest box size is an indication of thermodynamic favoring of T in the largest water box.

### Incorrect assertion about effect of iron-histidine bond on transition times

The response of El Hage et al. states, in relation to Fig 4, “Panels (A), (B), and (D) show that the presence or absence of the Fe-His bond does not lead to substantial changes of the time series as far as they have been followed.” Again, this statement is based on single, short simulations in which the bond between the heme iron and the proximal histidine was modeled by a covalent bond instead of by non-bonded interactions as in the original work (1). As before, we argue that single trajectory evidence is anecdotal and thus not statistically sufficient to support such a statement.

In addition to the statistical concern about the assertion that the presence or absence of the Fe-His bond does not lead to substantial changes of the transition times, we here show evidence that this claim is incorrect. As can be seen in Fig. 4, the introduction of a covalent bond to describe the interaction of the heme iron to the proximal histidine leads to significantly faster transition times.

### Unaltered conclusion

In the response by El Hage et al., the authors “argue that our original conclusions remain valid” (3). As posted at the beginning of this comment, all three of our original concerns remain, and none has been addressed in the response from El Hage et al. With the questioned issues on statistics, normalization and thermodynamics, that render the conclusions unsupported by the presented results, it is remarkable that the authors maintain their original conclusions. One of the main criteria in the assessment of a paper considered for publication is whether the conclusions are sufficiently backed up by the data. This appears not to be the case here.

### Technical concerns

El Hage et al. in their response (3) include a number of incorrect assertions about setup and analyses in our comment (2).

- “The subvolume analysis which attempts to compare the diffusivity in large boxes but restricted to a smaller subvolume needs to introduce ‘imaginary boundary conditions’ and it is unclear how this was done.“ No subvolume analysis was carried out for the diffusion constant analysis included in our comment.
- “A possible problem with the comparisons in Figure 3 of the comment concerns the fact that we only exchanged input files for the 9 nm box”. Here we’d like to point out that it was the choice of the authors to only send us the 9 nm box.
- “In that regard, it is interesting to note that their observed life time for the 9 nm box with its error bars essentially includes the one observed in our simulation.” Here we’d like to point out that this is also the case for the 12 nm and 15 nm boxes, therefore validating, rather than questioning, our setup for those box sizes as they behave indistuinguishably from those of El Hage et al. the protonation states in Table 1 as reported for the El Hage et al (2018) article do not match those of the actual simulations.
- “the terms for the dihedral parameters for the Fe-proximal histidine differed.” Firstly, it is important to underscore that our hemoglobin simulations labeled “El Hage” followed the El Hage et al. setup, using identical molecular topologies, including dihedral terms, to those reported in (1). Secondly, for the additional hemoglobin simulation setups labeled “Hub” and “Kovalevsky” a covalent Fe-proximal bond was used. The dihedral terms are a secondary issue to the presence or not of a covalent bond to model the iron interaction with the proximal histidine. It is common practice in the hemoglobin community to model this interaction as a covalent bond to represent the short iron-nitrogen distance. In the El Hage et al. setup this bond was absent, thus modeling both the iron interactions for the proximal as well as the distal histidine solely with non-bonded interactions.
- “Removing the angular center of mass motion is only warned against if the motion of the center of mass itself is not controlled, i.e. if the solute can cross the boundaries of the periodic box.” This is incorrect. Removing the angular center of mass motion in a periodic system is always warned against as it results in artifacts. Indeed, running with the El Hage et al. setup, where this problematic setting is switched on, GROMACS warnings are triggered that need to be manually overriden to be able to continue. Using this setup leads to severe issues with energy conservation.
- “the solute can cross the boundaries of the periodic box. This, however, is not the case in our simulations, but appeared to happen in some of theirs.” There are two flaws in this claim. Firstly, the solute moving across the periodic box boundaries is a concern only if angular center of mass motion removal is used. Without the center of mass removal, molecules, independent of being solvent or solute, are constantly moving across the periodic box boundaries in MD simulations without creating issues. Secondly, for the direct comparison of our hemoglobin simulations with those of El Hage et al., we used simulation parameters provided by El Hage et al. This way our trajectories were generated under identical conditions to those of El Hage et al.
- “Concerning the effect of the box shape on the proteins simulated under PBC, it has been found that the box type can have a statistically significant effect on the outcome of a simulation”. The claim in the El Hage et al. paper is not about box shape but about box size. In our comment we have explicitly tested the box size dependence for both cubic as well as rhombic dodecahedral box shapes and not found any significant effects for either. Whether or not the box shape affects the results in the hemoglobin case is an entirely different matter that is interesting to study by itself but is not the matter of debate here.
- “The results for the other two systems are interesting but there is no reason to expect a box size dependence for the structural changes that were studied.” The other systems that were studied (there were three additional systems, not two) we had included based on the final sentence of the abstract of the original El Hage et al. paper: “The possibility that extra large boxes are required to obtain meaningful results will have to be considered in evaluating existing and future simulations of a wide range of systems.” Here, no limitation to a specific class of systems or structural changes was made but instead effects for a wide range of systems were implied. However, in six out of six studied systems, no qualitative effects of the type claimed by El Hage et al were observed.
- “The role of the particular choice of His protonation states in the fast decay presented in the Comment remains to be evaluated.” The effect of histidine protonation on T->R transition kinetics for hemoglobin has already been evaluated in 2013 (7), which was however not cited by El Hage et al. (3).

## Conclusions

Our original concerns remain, as they were not addressed in the response of El Hage et al. In addition, the El Hage et al. response (3) gave rise to several additional concerns. Together with evidence based on additional data that we provide here, we conclude that no significant box size effects in MD simulations of hemoglobin have been demonstrated. Nature recently retracted a paper due to an issue with statistics (8), even though in that case the conclusions remained valid. In contrast, in the case of El Hage et al., which is more severe because the conclusions have been demonstrated to be unsupported, eLife has not pursued a retraction, neither of the original article (1), nor of the response comment (3). Moreover, with the publication of the response comment (3), eLife has endorsed re-stating of the original, unsupported conclusions and the introduction of further flawed supporting arguments. In addition, eLife has not taken any opportunity to publish any of the remaining or novel concerns, while at the same time urging others to watch out for statistical mistakes (9).

## Materials and Methods

We recomputed the solvent radial distribution function for the simulations of different box sizes previously published (2) (the simulations followed El Hage et al. setup (1)). In the current implementation we added the functionality to use a specified radius for normalization, rather than the default simulation box size. For the presented data in Fig. 2 we used a normalization distance of 4.25 nm radius. The code for this implementation is available under https://github.com/blauc/gromacs/tree/rdf.

The alanine dipeptide data was taken from previously published trajectories (2) and divided in 6000 trajectories of 1.7 ns each. The trajectory length was chosen such that on average, one transition takes place per sample, comparable to the hemoglobin simulations of 1 microsecond each. Samples of sizes 10, 200 and 500, as well as regular intervals in between, were selected randomly from the generated trajectories. Transition times were recorded as described previously (2) and the mean and standard error for each sample size was recorded.

The hemoglobin simulations depicted in Fig. 1 were taken from previous work (2). For the data presented in Fig. 4, 80 additional repeats were added to the previous batch using El Hage et al. (1) simulation parameters and setup to yield a total of 100 simulations of 1 microsecond each. For the simulations with a bond between the heme iron and the proximal histidine, a covalent bond was modeled between the iron and the nitrogen in epsilon position of the proximal histidine side chain. In addition to the covalent bond, also the corresponding bond angles and dihedral angles were added. All other simulation parameters were identical to the El Hage et al. setup.

## Acknowledgements

This work was supported by BioExcel CoE (www.bioexcel.eu), a project funded by the European Union contracts H2020-INFRAEDI-02-2018-823830 and H2020-EINFRA-2015-1-675728. We thank Christian Blau for the custom RDF implementation.

## References

1. Krystel El Hage, Florent Hedin, Prashant K Gupta, Markus Meuwly, and Martin Karplus. Valid molecular dynamics simulations of human hemoglobin require a surprisingly large box size. eLife, 7:e35560, jul 2018.

2. Vytautas Gapsys and Bert L de Groot. Comment on ‘valid molecular dynamics simulations of human hemoglobin require a surprisingly large box size’. eLife, 8:e44718, jun 2019.

3. Krystel El Hage, Florent Hedin, Prashant K Gupta, Markus Meuwly, and Martin Karplus. Response to comment on ‘valid molecular dynamics simulations of human hemoglobin require a surprisingly large box size’. eLife, 8:e45318, jun 2019.

4. Bernhard Knapp, Luis Ospina, and Charlotte M Deane. Avoiding false positive conclusions in molecular simulation: the importance of replicas. Journal of Chemical Theory and Computation, 14(12):6127–6138, 2018.

5. Massimo Pigliucci and Maarten Boudry. Prove it! the burden of proof game in science vs. pseudoscience disputes. Philosophia, 42(2):487–502, Jun 2014.

6. Xavier Daura, Bernhard Jaun, Dieter Seebach, Wilfred F. van Gunsteren, and Alan E. Mark. Reversible peptide folding in solution by molecular dynamics simulation. J. Mol. Biol., 280:925–932, 1998.

7. Martin D. Vesper and Bert L. de Groot. Collective dynamics underlying allosteric transitions in hemoglobin. PLOS Computational Biology, 9(9):1–8, 09 2013.

8. L. Resplandy, R. F. Keeling, Y. Eddebbar, M. K. Brooks, R. Wang, L. Bopp, M. C. Long, J. P. Dunne, W. Koeve, and A. Oschlies. Retraction note: Quantification of ocean heat uptake from changes in atmospheric o2 and co2 composition. Nature, 573:614, 2019.

9. Tamar R Makin and Jean-Jacques Orban de Xivry. Science forum: Ten common statistical mistakes to watch out for when writing or reviewing a manuscript. eLife, 8:e48175, oct 2019.

